# Evaluating tank acclimation and trial length for shuttle box temperature preference assays

**DOI:** 10.1101/2020.07.21.214080

**Authors:** Adam Alexander Harman, Meghan Fuzzen, Lisa Stoa, Douglas Boreham, Richard Manzon, Christopher M. Somers, Joanna Yvonne Wilson

## Abstract

Thermal preferenda are largely defined by optimal growth temperature for a species and describe the range of temperatures an organism will occupy when given a choice. Assays for thermal preferenda require at least 24 hours, which includes a long acclimation to the tank, limits throughput and thus impacts replication in the study. Three different behavioral assay experimental designs were tested to determine the effect of tank acclimation and trial length (12:12, 0:12, 2:2; hours of tank acclimation: behavioral trial) on the temperature preference of juvenile lake whitefish, using a shuttle box system. Average temperature preferences for the 12:12, 0:12, and 2:2 experimental designs were 16.10 ± 1.07 °C, 16.02 ± 1.56 °C, 16.12 ± 1.59°C respectively, with no significant differences between the experimental designs (p= 0.9337). Ultimately, length of acclimation time and trial length had no significant impact, suggesting that all designs were equally useful for studies of temperature preference.

## Introduction

Most motile species are thought to exhibit a thermal preferenda or a range of preferred temperatures that individuals will tend to aggregate at when given the opportunity (Reynolds and Casterlin, 1979). This temperature should theoretically correlate with the optimum growth temperature, but there are several other important factors contributing to a thermal preferenda, including photoperiod, salinity, chemical exposure, age and/or size of fish, bacterial infection, nutritional state/food availability, and other biotic factors (Reynolds and Casterlin, 1979).

The definition of final preferenda assumes a common temperature preference that all members of the same species will ultimately display (Jobling, 1981). This may be accurate for small warm-water fish, like goldfish (*Carassius auratus*) and bluegill sunfish (*Lepomis macrochirus*), that were used for much of the early preferenda work (Reynolds and Casterlin, 1979) because they experience warm, stable temperatures across their distribution. The same cannot be said for larger temperate species that have consistently dealt with extreme temperature changes over their evolutionary history. Atlantic cod (*Gadus morhua*) display significantly different preferenda across their distribution due to a polymorphic haemoglobin molecule (Petersen and Stefensen, 2002), while juvenile coho salmon (*Oncorhynchus kisutch*) have distinct thermal preferences that align with the thermal profile of home streams (Konecki, 1995). Arctic charr (*Salvelinus alpinus*) that are exposed to repeated freezing and thawing of lakes/streams, experience seasonal changes in preferenda (Mortensen et al., 2007).

Temperature preference (T_pref_) in juvenile lake whitefish (*Coregonus clupeaformis*) is inversely related to the size and age of the fish (Edsall, 1999), suggesting that conspecifics of different age classes may show different temperature preferences within the same body of water. Further, the basal metabolic rate of a fish has been correlated to their aerobic scope and their temperature preference (Killen et al. 2014). Fish with higher basal metabolic rate have both a lower aerobic scope and temperature preference. To compensate for increased metabolic demands, fish with higher basal metabolic rate tend to select colder temperatures when food availability is low (Killen et al., 2014). Therefore, individual life history traits can account for differences in T_pref_.

Thermal preferenda assays are conducted in tanks with either a temperature gradient or a choice between different temperatures. These assays require an initial tank acclimation period where fish acclimate to the test arena, followed by a behavioral trial. Traditionally, the total assay (acclimation and trial) have a minimum length of 24 hours (Mortensen et al., 2007; Siikavoupio et al., 2014; Konecki et al., 1995; Petersen and Stefensen, 2002), based on the theory that fish are only displaying their acute temperature preference, rather than their final preferenda, when <24 hours in a new system (Reynolds and Casterlin, 1979). Allowing the fish to remain in the new system for at least 24 hours would theoretically reveal their final preferenda. However, Macnaughton et al. (2018) determined that tank acclimation time had little effect on the final preferenda of juvenile cutthroat trout (*Oncorhynchus clarkia lewisi*), a cold-adapted fresh-water species. Further, a minimum 24-hour assay length per fish has significant disadvantages for sample size and throughput in any study. The ability to assess preferenda would be extremely challenging in experiments that focus on biotic and abotic influences and fast growing life stages because of issues (e.g. length of time for experimental treatment, time out of treatment during the assay, different body sizes) inherent with the total time needed if throughput is ≤ 1 fish per day.

Fish in the juvenile life-stages, including lake whitefish, are in a period of rapid development and growth (Rennie, 2009), and Edsall (1999) reported a relationship between size and temperature preference. Long assay lengths may correspondingly introduce growth as a confounding factor. The influence on preference from seasons, migration, or physiological transitions with small temporal windows (e.g. smoltification), are difficult to determine because of limited throughput. Consequently, many studies (Mortensen, 2007; Barker et al., 2018; Larsson 2005; Petersen and Stefensen, 2002; Siikavuopio, 2014) use low sample sizes and have low statistical power. Alternatively, some studies test multiple fish at one time (Edsall, 1999; Sauter et al., 2001) but the social context likely influences results and individual fish are not truly independent measures. Increasing throughput would have significant advantages for all of these scenarios.

A shuttle box, first described by Neill (1972), is an instrument that determines the temperature preference of aquatic animals by allowing them to choose between two tanks held at different temperatures. Once acclimated to the system, fish will ‘shuttle’ between the two compartments to regulate body temperature, allowing analysis of preferred temperature and avoidance temperatures. This study examined the effect of tank acclimation and trial length on the quality and quantity of data produced to determine thermal preference (T_pref_) during behavioral assays. We used three distinct experimental designs, starting with a 24-hour total assay length (12 hours tank acclimation:12 hours trial length) as a baseline. It was hypothesized that experimental designs of different lengths (24 hours, 12 hours, 4 hours) would have a limited effect on the determined thermal preference of lake whitefish (*Coregonus clupeaformis*) and that shorter assay designs could increase throughput.

## Methods

Fertilized lake whitefish (LWF) embryos were acquired from Sharbot Lake White Fish Culture Station (Sharbot Lake, ON) on November 30^th^, 2017. Embryos were incubated under simulated seasonal temperatures until hatch. Embryos were initially held at 8°C and cooled (1°C/week) to 2°C. After 100 days of incubation, embryos were warmed (1°C/week) until hatching. Median hatch occurred at 158 days post fertilization. Hatchlings were placed in petri dishes at 8°C until successful exogenous feeding. Larvae were transferred to tanks and warmed (1°C/week) to 15°C, where they remained until testing (5-6 months). LWF were initially fed *Artemia* nauplii and slowly transitioned to pellet feed (Otohime B1 (200-360 μm) – C2 (920-1,410 μm) larval feed).

The shuttle box system (Loligo^®^) consists of two cylindrical tanks connected by a small rectangular ‘shuttle’ to allow movement of animals between the tanks. Each tank is assigned as the increasing (INCR) or decreasing (DECR) side, indicating the direction of temperature change when fish occupy that tank. To accurately regulate temperature, system water was pumped through heat-exchange coils in hot (28°C) and cold (4°C) water baths (60L aquaria) with mixing in separate buffer tanks for each side. A Recirculator 1/4 HP Chiller, Magnetic Drive Centrifugal Pump (300W/600W/950W @ 0°C/10°C/20°C; VWR) and a 400W aquarium heater were used to maintain the temperatures in the cold and warm bath, respectively. Ice was added to the cold bath every 2 hours during shuttle box operation to increase cooling capacity. Polystyrene insulation (1/2″), foam insulation tape (1/4″), and loose fiberglass insulation were used to maintain stable temperatures in the cold-water bath. System water flows (240 mL/min) via gravity through temperature probes and into the shuttle box where counter-directional currents minimize mixing between the two sides. A USB 2.0 uEye Camera tracked larval fish under infrared light (Loligo^®^ Infrared Light Tray), and the Shuttlesoft^®^ software determined the ‘live’ location of the tracked object. Shuttlesoft^®^ uses contrast to identify and track objects and required even, symmetrical overhead lighting; black opaque plastic was used to dim fluorescent lights directly overhead and prevent glare.

In our experiments, we defined distinct static or dynamic modes for the shuttle box; the total assay length was the sum of time for each mode. Static mode (tank acclimation) was used to acclimate the fish to the shuttle box system but was not used to determine temperature preference. In this mode, the shuttle box maintained stable temperatures of 14°C and 16°C with a hysteresis of 0.25°C. Dynamic mode (behavioral trial) was used to determine temperature preference; fish were actively tracked and the entire system would warm or cool (hysteresis = 0.1°C) at a rate of 4°C/hour, depending on whether the fish was in the INCR or DECR tank. In both static and dynamic modes, the difference in temperature across the tanks was Δ 2°C. Hysteresis values were determined experimentally for each operating mode independently to achieve the most stable water temperatures over time. A maximum temperature of 23°C and a minimum temperature of 7°C prevented exposure to extreme temperatures, which could cause stress or mortality (Edsall and Rottiers, 1976).

The orientation of the INCR and DECR tanks and the side to which the fish would be introduced were randomized for each individual, using an online tool (random.org), to limit any potential bias introduced by visual cues or side preference. LWF were randomly selected from their home tank (15°C) and transported to the shuttle box system in 1L glass beakers. LWF were introduced to one side of the shuttle box, with a plastic divider separating the two halves. The assay started immediately after the barrier was removed, initiating acclimation, and continued until the end of the behavioral trial. While data were collected throughout, only data collected during the behavioral trial (dynamic mode) were used for temperature preference analysis. Shuttlesoft^®^ calculates temperature preference (T_pref_) over time as the median occupied temperature; velocity (cm/s), distance (cm), time spent in INCR/DECR, number of passages and avoidance temperatures were collected in 1 second intervals. The fish remained in the shuttle box throughout the entire assay, without interference or handling. After completion of the assay, fish were removed and measured for total length (±1 mm) and mass (±0.01 g) before returning fish to a separate home tank (15°C).

Three experiments were conducted to test the effect of tank acclimation and trial length on the quality of data, namely 12:12, 0:12, or 2:2 designs representing the number of hours in static mode (tank acclimation) and dynamic mode (behavioral trial), respectively (Figure 1a). Summary statistics were generated for each experimental design to compare the effect of the design on data accuracy and variability. Mean T_pref_ + standard deviation was used to compare the variation between fish, which is the major limit of statistical power. An experimental design was considered equally useful if it produced T_pref_ data that were not statistically different. Power analyses were completed for each experimental design to compare optimal sample sizes at the lowest acceptable power (1−β =0.60).

**Figure 1:**
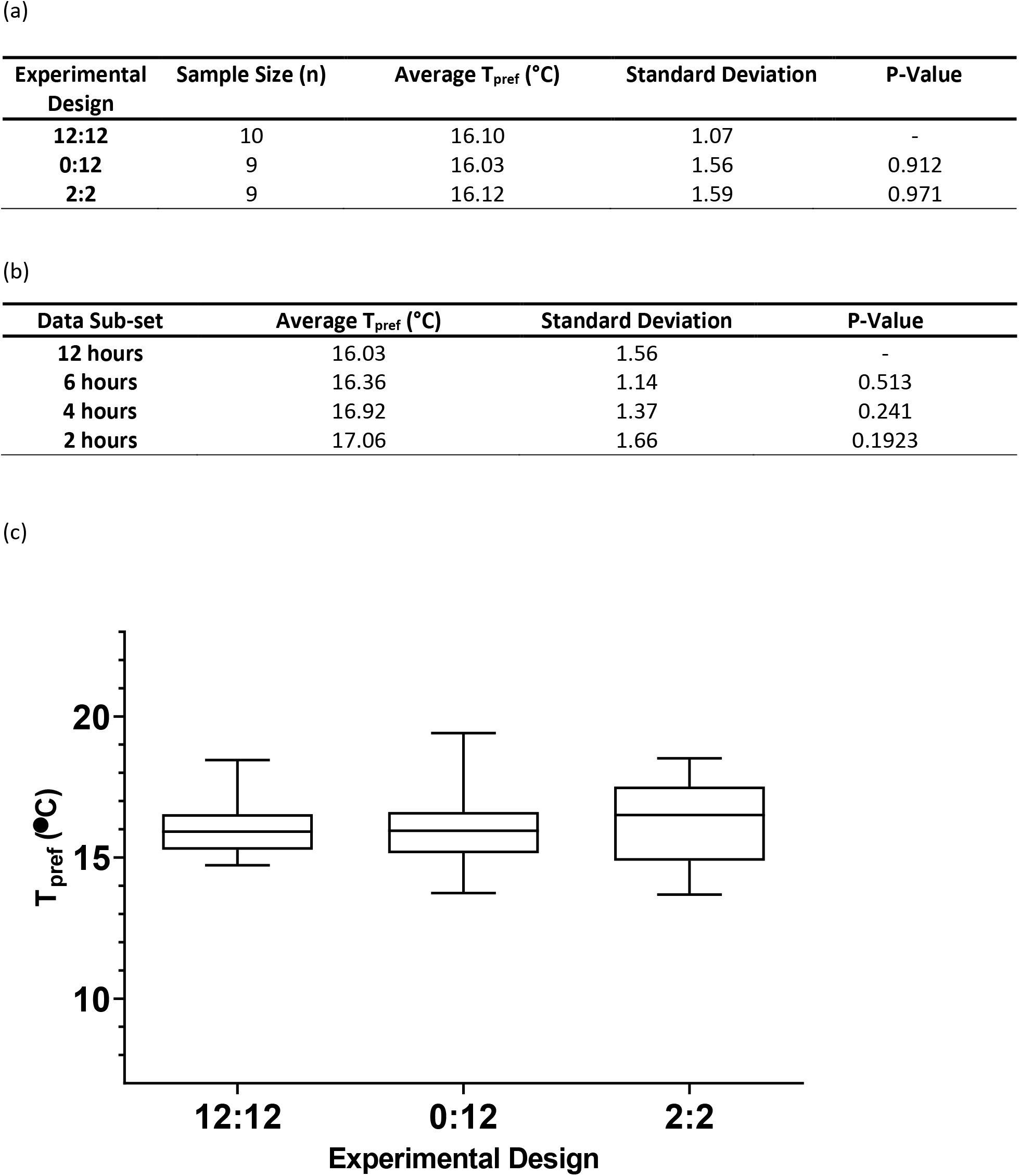
(a) Summary of average temperature preference (T_pref_) data from three different experimental designs. T_pref_ is calculated as the cumulative median of occupied temperature. 12:12, 0:12, or 2:2 designs representing the number of hours in static mode (tank acclimation) and dynamic mode (behavioral trial), respectively. P-values were determined using one way ANOVA with post-hoc comparisons. (b) Sub-set analysis conducted using the 0:12 experimental design, behavioral trials were sub-set into 2, 4, and 6-hour windows. P-values were determined using ANOVA. (c) Box plot comparing T_pref_ between 12:12, 0:12 and 2:2 experimental designs. The height of the box corresponds to Q1 – Q3, and the bars correspond to the minimum and maximum values. Y-axis represents the thermal range of the shuttle box system.

## Results and Discussion

In the first experimental design (12:12), juvenile LWF (n=10) had 12 hours of over-night tank acclimation (9 pm – 9 am) in static mode, followed by 12 hours of behavioral trials (9 am – 9 pm) in dynamic mode. The maximum throughput was 1 fish per day (Figure 2e). This design included the longest tank acclimation period, the lowest throughput and was predicted to decrease between-fish variability. The average T_pref_ was 16.10 ± 1.07 °C (Figure 1a), which was the lowest standard deviation in average T_pref_ across the experimental designs, as expected.

**Figure 2:**
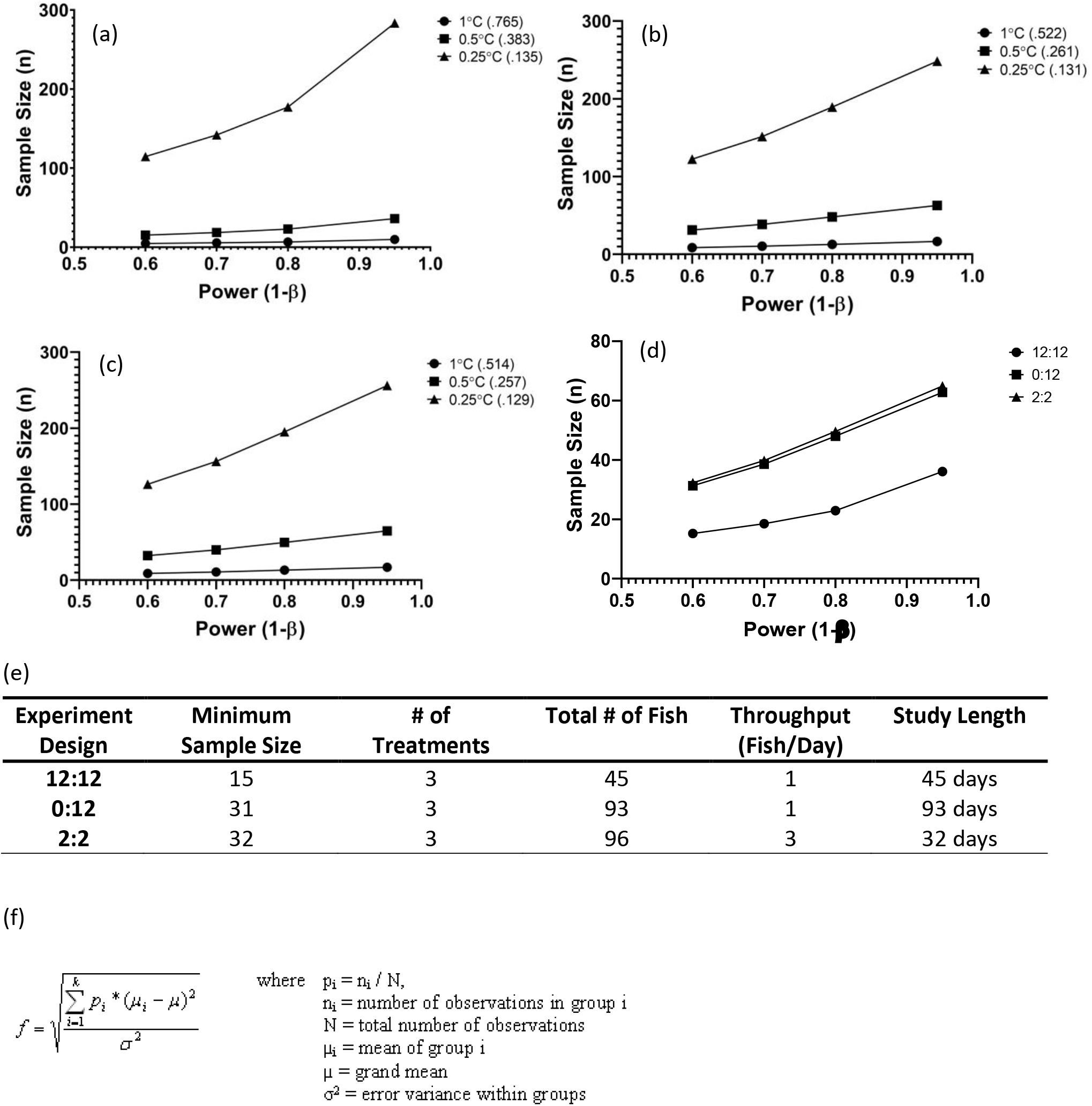
(a, b, c) Relationship between sample size (n) and power (1-β) for experimental (Expt) designs 12:12 (a), 0:12 (b), and 2:2 (c), representing the number of hours in static mode (tank acclimation) and dynamic mode (behavioral trial), respectively. Curves were generated using iterative power analysis (pwr package – R). Effect sizes were calculated using panel (f) by predicting expected differences between means. (d) Power analysis using 0.5°C effect sizes, each data series corresponds to an experimental design. (e) Summary of power analysis results. Minimum sample size corresponds to n calculated with 0.5°C effect size and 1-β = 0.6. # of treatments can vary with experimental design, three was chosen as a reasonable example. Total number of fish is minimum sample size times the number of treatments. Study length was calculated by dividing the total number of fish by the throughput of the experimental design, 12:12 = 1/day, 0:12 = 1/day, 2:2 = 3/day. (f) Equation used to calculate effect size (ƒ) for ANOVA.

Available literature suggests that a long tank acclimation period prior to the behavioral trial is required to observe the true temperature preference of a species (Reynolds and Casterlin, 1979). The second design (0:12) explicitly tested the effect of tank acclimation by completely removing it; juvenile LWF (n=9) had a 12-hour behavioral trial (9 am – 9 pm) under dynamic mode with no prior acclimation. One fish was excluded because the system shut down prematurely. Removal of the static period was predicted to increase the variation in T_pref_ between individuals. As predicted, the standard deviation of T_pref_ increased, but not drastically (Figure 1a). Throughput (1 fish/day) remained the same because only the overnight tank acclimation was removed; while 2 fish/day were possible if we ran assays in both day and night, results were more comparable with dynamic mode in the same part of the diurnal cycle (day light). The average T_pref_ was 16.03 ± 1.56 °C (Figure 1a), which was not statistically different (p=0.912) from the outcome using the baseline design. The data from this experiment were analyzed in 2-hour sub-sets (i.e. 2 hours, 4 hours, 6 hours) to simulate shorter behavioral trial durations (Figure 1b). Average T_pref_ was not statistically different (p=0.1923) between a 12-hour and a 2-hour behavioral trial length (Figure 1b), suggesting that not only was long tank acclimation not required but shorter trials were possible. The advantage of no or limited tank acclimation coupled with a shorter behavioral trial was that throughput could be increased to multiple fish per day, offering the opportunity to increase total sample size or decrease the time needed to assess T_pref_ in different treatment groups.

A third experimental design (2:2) was implemented with 2 hours of tank acclimation and 2 hours of behavioral trial, to increase throughput. Three time periods were used (11 am – 1 pm, 3 pm – 5 pm, 7 pm – 9 pm) instead of one (9 am – 9 pm), which would triple throughput; there was no effect of time of day. This design has not been reported in the literature and this is the first attempt to calculate T_pref_ from such a short assay, to our knowledge. The average T_pref_ was 16.12 ± 1.59°C (Figure 1a) and was not significantly different from either alternative experimental design (p=0.9337). Further, the standard deviation did not drastically increase (Figure 1a), although it was the largest of the tested designs.

Shuttlesoft^®^ automatically calculates the cumulative median of T_pref_ every second, and that data can be compared between individuals and groups. Figure 3 compares individual T_pref_ data to the average, showing the spread of the data as well as the stability over time. A unique aspect of the shuttle box behavioral assay is that a fish must be shuttling between the two sides to maintain a constant temperature within the system; switching sides is an active behavioral choice. Traditional methods require the fish to remain stationary to select a temperature in a gradient. All experimental designs followed a similar pattern of an initial period of high variability, followed by a prolonged period of relative stability (Figure 3), suggesting an active choice was made. Therefore, the different designs appear largely equivalent, suggesting that long tank acclimation and long behavioral trials are not necessary to determine T_pref_, at least for juvenile LWF. This offers the opportunity to increase the throughput on a temperature preference study where confounding variables (e.g. rapid body growth, exposure to abiotic or biotic factors) could significantly impact the data if the traditional design (>24 hours per fish) was used.

**Figure 3:**
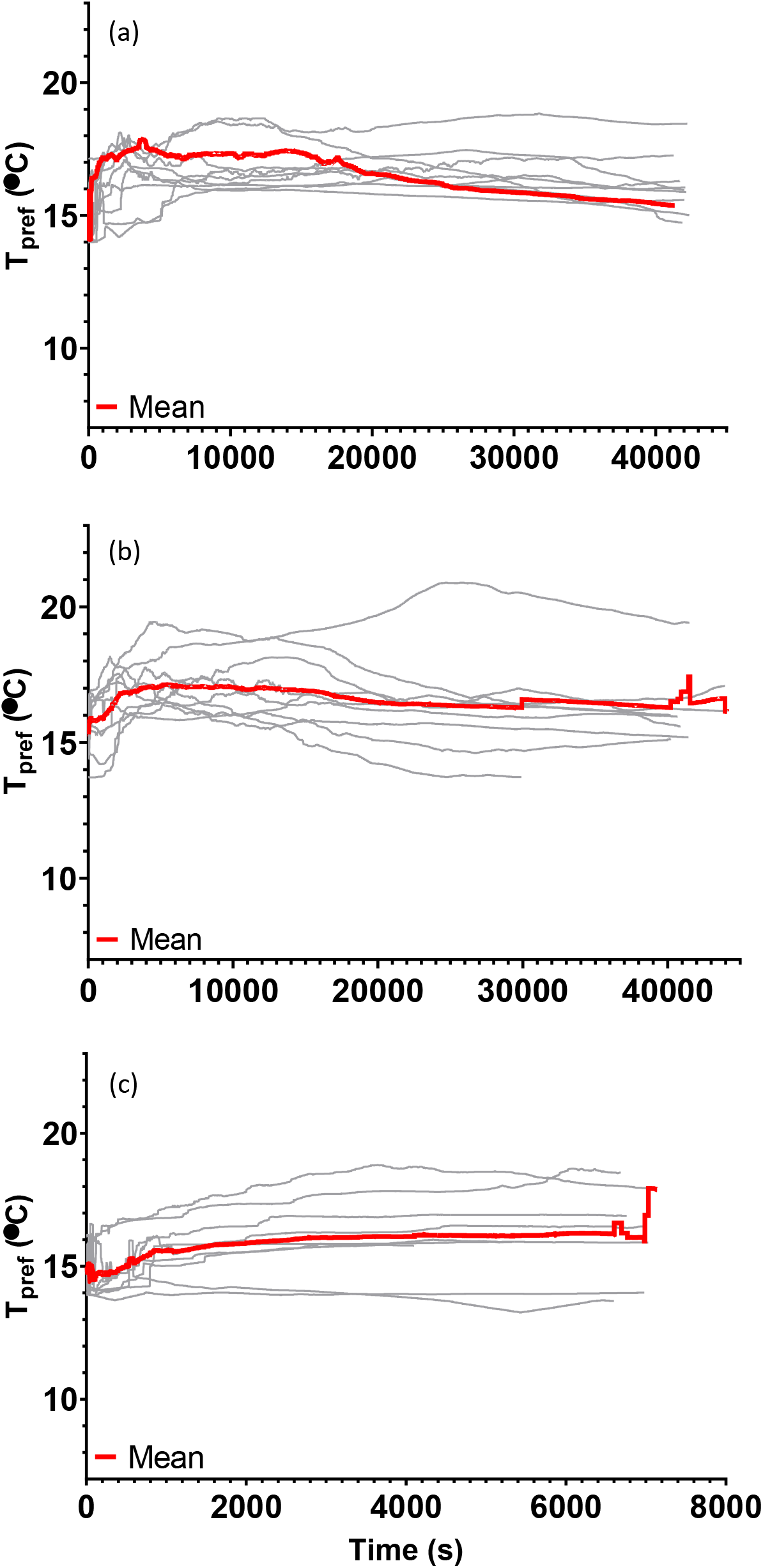
Cumulative median temperature preference (T_pref_) calculated every 1 second for experimental designs 12:12 (a), 0:12 (b) and 2:2 (c), representing the number of hours in static mode (tank acclimation) and dynamic mode (behavioral trial), respectively. Grey lines represent the T_pref_ of individual fish over time. Red line represents the mean T_pref_ for all fish. Y-axis represents the thermal range of the shuttle box system.

Tank acclimation and behavioral trial intervals were chosen based on both scientific evidence and logistics. In all cases, we note the throughput (i.e. how many fish can be tested per week) to highlight the relevant trade off that would impact experimental design choice. While previous literature (Mortensen et al., 2007; Siikavoupio et al., 2014; Konecki et al., 1995; Petersen and Stefensen, 2002) would suggest acclimating fish to the tank for a period of >24 hours, we used a total assay length of 24 hours (12-hour static tank acclimation, 12-hour dynamic behavioral trial) as the baseline. This was chosen because a total assay length of >24 hours would lead to a throughput of only 3 fish/week, which would not have been feasible for a large-scale experiment, particularly with fast growing juvenile fish. Considering the juvenile fish used here (5 months of age), it would be important to account for changes in individual growth during temperature preference studies. A negative correlation between growth and temperature preference has been observed in lake whitefish (Edsall, 1999), which suggests study length could be an influential factor in experiments with fast growing life stages. Increasing throughput could allow testing a wider range of individuals (Figure 2e) and may better capture a population’s natural variability.

Using the 2:2 design would yield an experiment that is 34 days in length to provide the minimum sample size needed for three treatment groups (Figure 2e). Even within 34 days, individual juvenile LWF tested near the beginning of the study would be ~20% younger and 11% smaller (LWF are 9.11 g (± 2.8) versus 10.23 g (± 2.0) at 5 and 6 months, respectively; unpublished data). It would be important to minimize length of time to collect temperature preferenda data and consider the trade-offs between variance and sample size on the statistical power to assess differences across treatment groups. The same can be said when determining T_pref_ within small temporal windows (e.g. smoltification, seasonality, developmental windows) where small sample sizes would limit statistical power. The functional trade-offs between statistical power (1-ß), variance (δ^2^), sample size (n), and throughput were investigated using power analysis (Figure 2) for the various experimental designs. While experimental design 3 (2:2) led to increased variation in mean T_pref_, the increased throughput allowed for an increased sample size while still minimizing the total time needed for the experiment. If the number of fish were limited or growth and developmental concerns were not as relevant (e.g. adult fish), then minimizing variation may be more important.

This study used a maximum rate of change of 4 °C/hour, similar to what has been previously reported (Macnaughton et al., 2018; Konecki, 1995; Petersen and Stefensen, 2002). This could have limited the range of temperatures experienced by the juvenile LWF. If a fish occupied the INCR zone for the entire duration of the behavioral trial, the system would have cooled by 8°C, only just hitting the upper temperature limit of the shuttle box. Thus, to reach extreme temperature preferences a fish must exhibit low (<10) passage numbers, a problem when preference is determined by active swimming. This problem could potentially be avoided by increasing the rate of temperature change (Barker et al., 2018), at the expense of possible physical stress. For our experiments, data were excluded only when fish made no passages in the dynamic mode. In all cases, fish made regular passages in at least one mode, indicating they were active and able to explore the entire arena. Hyperactive fish would likewise pose a problem for the system; there was no animal that exhibited so many crosses that the system could not respond and change temperature.

Thermal preferenda can be an important behavioral endpoint but traditionally require long periods of time (>24 hours) to determine. The results of this study show that decreasing the total assay length (24 hours to 4 hours) did not significantly affect the T_pref_ of juvenile lake whitefish. The shuttle box is a powerful behavioral tool and a less restrictive definition of T_pref_ and more flexibility in the assay design would allow T_pref_ as a viable behavioral endpoint for a variety of species and life stages with more experimental power.

